# Mosaic-PICASSO: accurate crosstalk removal for multiplex fluorescence imaging

**DOI:** 10.1101/2023.07.06.547878

**Authors:** Hu Cang, Yang Liu, Jianhua Xing

## Abstract

Ultra-multiplexed fluorescence imaging has revolutionized our understanding of biological systems, enabling the simultaneous visualization and quantification of multiple targets within biological specimens. A recent breakthrough in this field is PICASSO, a mutual-information-based technique capable of demixing up to 15 fluorophores without their spectra, thereby significantly simplifying the application of ultra-multiplexed fluorescence imaging. However, this study has identified a limitation of mutual information-based techniques. They do not differentiate between spatial colocalization and spectral mixing. Consequently, mutual information-based demixing may incorrectly interpret spatially co-localized targets as non-colocalized, leading to overcorrection. We found that selecting regions within a multiplex image with low spatial similarity for measuring spectroscopic mixing results in more accurate demixing. This method effectively minimizes overcorrections and promises to accelerate the broader adoption of ultra-multiplex imaging.

## Introduction

The introduction of multiplexed fluorescence imaging methodologies, such as CODEX^1^, MIBI^2^, CyCIF^3^, 3i^4^, CosMX^5^, and ImmunoSABER^6^, has revolutionized biological and medical research by providing unprecedented capability to visualize molecular complexities. However, the challenge of spectral overlap limits the concurrent imaging of fluorophores to a maximum of four^7^. This constraint necessitates cyclic multi-round imaging, which is laborious and susceptible to morphological disturbances. Linear unmixing is an appealing solution to spectral overlap that has been proposed. However, the procedure requires accurate quantification of a mixing matrix, which has to be calibrated every time in each experiment, thus adds complexity.

Recently, a groundbreaking approach, PICASSO (Process of ultra-multiplexed Imaging of biomoleCules viA the unmixing of the Signals of Spectrally Overlapping fluorophores)^7, 8^ has been developed. It is capable of unmixing up to 15 colors in a single imaging round. Notably, PICASSO employs mutual information (MI) of images from different channels to quantify the spectroscopic mixing of diverse fluorophores. It does not require any prior knowledge of the reference fluorescence spectra of the fluorophores. This blind unmixing approach significantly alleviates the burdens for users to generate multiplex fluorescence imaging, circumvents errors in reference measurements, and has the potential to revolutionize the field of multiplex imaging.

The framework of PICASSO is based on the assumption that mutual information is an indicator of spectroscopic mixing. By minimizing the mutual information, the mixing parameters can be estimated, thus facilitating successful unmixing. However, there are scenarios where mutual information between two color channels does not reflect the true biological processes, such as co-localizations between two proteins. Indeed, we found that the minimization of mutual information from PICASSO’s spectroscopic mixing model can erroneously interpret the spatially co-localized targets as spectroscopically mixed signals between two color channels, leading to overcorrection. Consequently, the demixed images portray spatially co-localized targets as spatially non-colocalized, potentially impacting the accuracy of the unmixing process.

To address this limitation, we propose to harness the spatial heterogeneity within a biological sample. An image can be divided into regions with low and high spatial similarities. By selectively applying PICASSO to areas of low spatial similarity, we can better estimate the mixing parameters compared to full-image measurements that encompass regions with both low and high similarity. Subsequently, we utilize the measured mixing parameters to demix the entire image. This mosaic-based approach preserves the inherent blind demixing capability of PICASSO while effectively overcoming its limitations in highly spatially overlapped samples. The utilization of mosaic-based PICASSO significantly enhances the accuracy of the unmixing process, resulting in a substantial reduction in errors in the demixed images.

## Results

### PICCASO achieves spectroscopic demixing through minimization of mutual information

Mutual information of two images quantifies the amount of information one can derive about one image based on the other, assessing the statistical interdependence between the intensities of the two. For channels Img1 and Img2 of the same image, the normalized Mutual Information (MI) is calculated using the following formula:

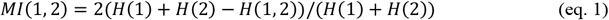

Here, H(1) is the entropy of channel 1, H(2) is the entropy of channel 2, and H(1, 2) is the joint entropy of channels 1 and 2. The entropy H of an image could be calculated from the pixel intensity. The normalized mutual information is in the range of 0 (no correlation) and 1 (exactly the same). When mutual information is high, it implies a strong alignment between the pixel intensities of the two channels.

Mutual information has been successfully deployed in the past for registering images from different modalities^9^. PICASSO innovatively applies the principle of mutual information minimization for spectral unmixing, operating under the assumption that spectral mixing enhances the mutual information between multiple channels. Consequently, the unmixed images can be retrieved by minimizing this mutual information.

For simplicity, let’s first consider a two-color image with Img1 and Img2 representing the observed images from two fluorescent channels, respectively (Fig. 1a). Fluorophores 1 and 2, which primarily emit in Img1 and Img2, exhibit leakage into the other channels. Consequently, the measured images Img1 and Img2 contain spectral leakage from these fluorophores, thus they are mixed. That is, Img1 and Img2 are related to the actual images F1 and F2 through linear combinations as follows:

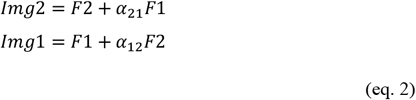

**Fig. 1:**
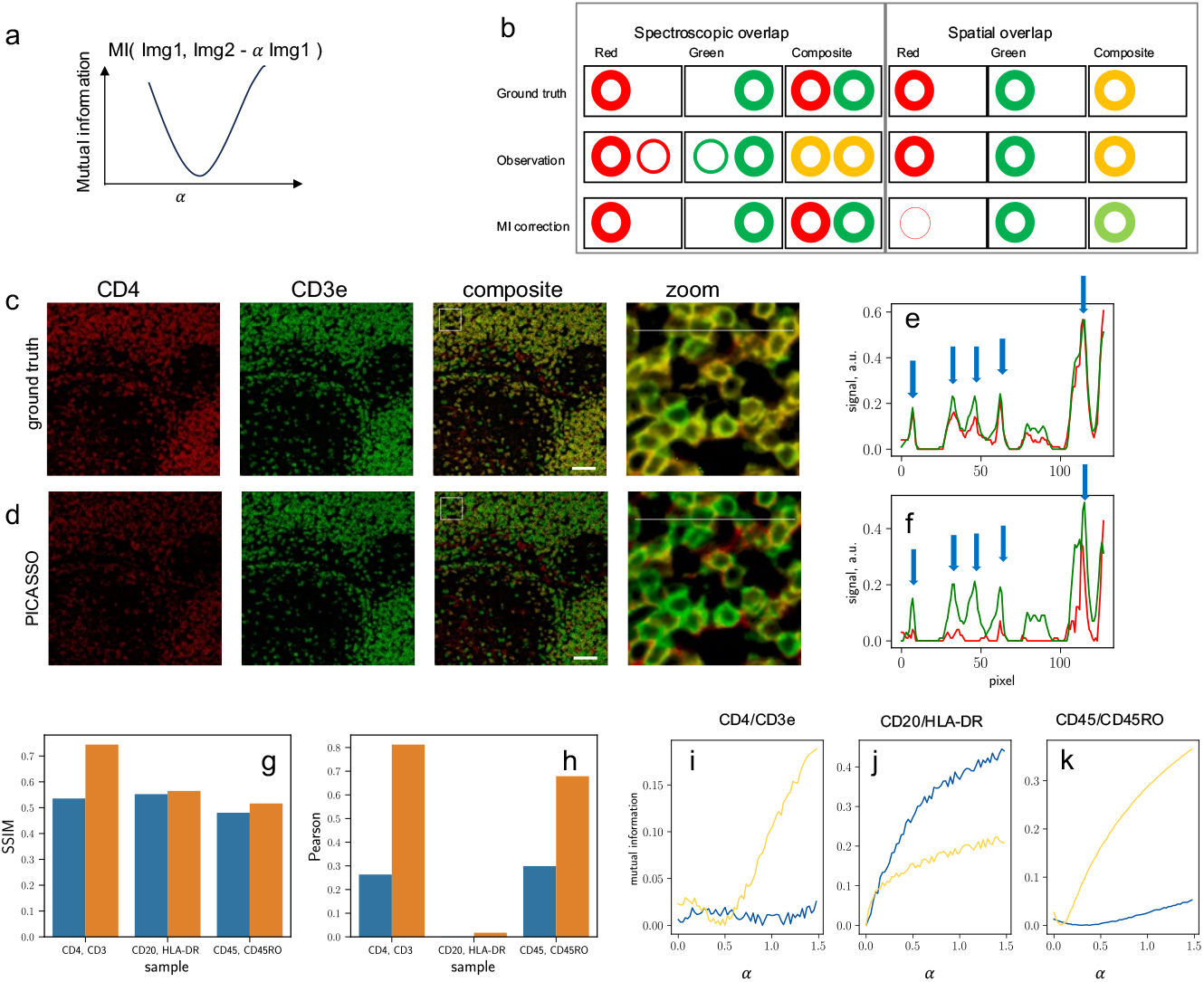
Limitations of using mutual information in resolving spectroscopic and spatial overlaps. (a) Illustration of the algorithm for eliminating spectroscopic cross-talk using mutual information. (b) Increase in mutual information can be due to both spectroscopic overlapping and spatial co-localization of fluorophores. Minimizing mutual information may mistakenly classify co-localized fluorophores as non-colocalized. (c, d) Over-corrections induced by PICASSO on spatially co-localized fluorophores. (e, f) Profiles of the two channels. Blue arrows indicate discrepancies between ground truth and PICASSO results. (g) SSIM structural similarity index of ground truth (yellow) and PICASSO-corrected 2-channel images (blue). Samples with lower overlap (CD20/HLA-DR) and (CD45/CD45RO) exhibit less overcorrection compared to highly overlapped samples (CD4/CD3). (h) Pearson’s correlation reveals similar overcorrection artifacts for highly overlapped sample (CD4/CD3). (i-k) MI curves plotted against the mixing ratios *α*_12_ (blue) and *α*_21_ (yellow). The blue and yellow curves are calculated as MI(red, green - *α*_12_ red) and MI(green, red - *α*_21_ green), respectively. The estimated mixing ratios are obtained from the minimal points of the curves.

Here, to effectively unmix the color channels and reduce the dependency between them, the PICASSO approach searches for the values of α_12_ and α_21_ that minimize the mutual information between Img1 and Img2, such that:

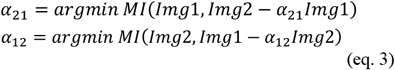

The PICASSO algorithm can be applied to an arbitrary number of channels using an iterative optimization process^7^. In each step, PICASSO selects a pair from multi-channel images; it then estimates the level of mixing between the two channels by (eq. 3). PICASSO then subtracts the scaled images from other images and repeats this process for all possible pairs. This procedure is iteratively performed to minimize the mutual information.

### Spatial co-localization confounds the performance of PICASSO

However, mutual information-based spectral unmixing methods cannot differentiate between spectroscopic mixing and co-localization of fluorophores. This problem is especially severe for those images at low-to-moderate resolution, where many fluorophores can be spatially co-localized. Fig. 1b illustrates this limitation, where we consider the images of cells with two different surface markers (red and green). In the case where the cells are spatially separated but spectrally mixed (left panel), the composite image appears as two yellow circles due to the spectral leakage from the other channel. By minimizing mutual information, the leakage can be perfectly removed, resulting in the unmixing of the images into a red and a green circle. However, when the two fluorophores co-localize on the same cells, the composite image also appears as yellow, similar to the spectral mixing scenario. Minimizing mutual information in this case erroneously removes the co-localization, resulting in the composite image, which should be yellow, appearing as a single color.

To test the impact of co-localization on spectral unmixing, we used images taken by CyCIF^3^ (Fig. 1c-f) for a test. In our analysis, we focused on the CD4 and CD3 targets. The images were acquired over multiple cycles, with each cycle involving cleavage of the previously used fluorophore, resulting in minimal cross-talk or mixing between the two channels. Therefore, the raw images can be considered as ground truth with no spectral mixing.

However, PICASSO reports significant spectroscopic mixing between CD4 and CD3, resulting in substantial subtraction of one of the channels during the unmixing process. The cross-section plot of the ground truth image (Fig. 1e) clearly demonstrates the near-perfect co-localization of CD4 and CD3 on the cell surface. However, after the unmixing, they become completely de-correlated, suggesting the potential risk of overcorrection in spectral unmixing algorithms (Fig. 1f). We have also tested other targets, CD20/HLA-DR and CD45/CD45RO, which exhibit a lower degree of spatial colocalization. Additional data showcasing overcorrections of highly spatially overlapped targets can be found in the supplementary figures (Supplementary Fig. 1 and Supplementary Fig. 2).

To further evaluate the impact of the demixing process, we employed commonly used colocalization metrics, namely the Pearson correlation coefficient and the Structural Similarity Index (SSIM), to measure the similarity between different colors (Fig. 1g-h). Both the SSIM (Fig. 1g) and the Pearson correlation coefficient (Fig. 1h) showed a significant decrease after demixing, indicating substantial mis quantification of the co-localizations.

Fig. 1i-k illustrate the mutual information curve for the three images (CD4/CD3e, CD20/HLA-DR, and CD45/CD45RO) plotted against the mixing ratios *α*_12_ and *α*_21_. The minimal point of the mutual information curve represents the estimated mixing ratios. In contrast to the expected ground truth values of zero for all samples, the presence of spatial colocalization results in positive mixing ratios when minimizing mutual information. This positive mixing manifests as over-subtraction. Moreover, the degree of overcorrection varies depending on the level of spatial colocalization. Highly overlapped targets (CD4/CD3e) exhibit more significant overcorrection compared to lower overlapped targets (CD45/CD45RO and CD20/HLA-DR). These findings highlight the complexity in interpreting the biological implications of the unmixed images.

### Spatially heterogeneous unmixing results

We observed that the results of the unmixing process can exhibit high spatial heterogeneity depending on the selection of regions of interest (ROIs) within an image. To investigate this, we divided the CD4/CD3e image into small mosaics (Fig. 2a) and performed independent PICASSO analysis on each mosaic. The size of the mosaics was chosen to ensure that each mosaic contains typically one to two cells on average. This selection allows for a fine-grained analysis and assessment of spatial heterogeneity at a cellular level. Additionally, we calculated the Structural Similarity Index (SSIM) for each mosaic to assess the similarity between the two channels of each mosaic (Fig. 2b).

**Fig. 2.**
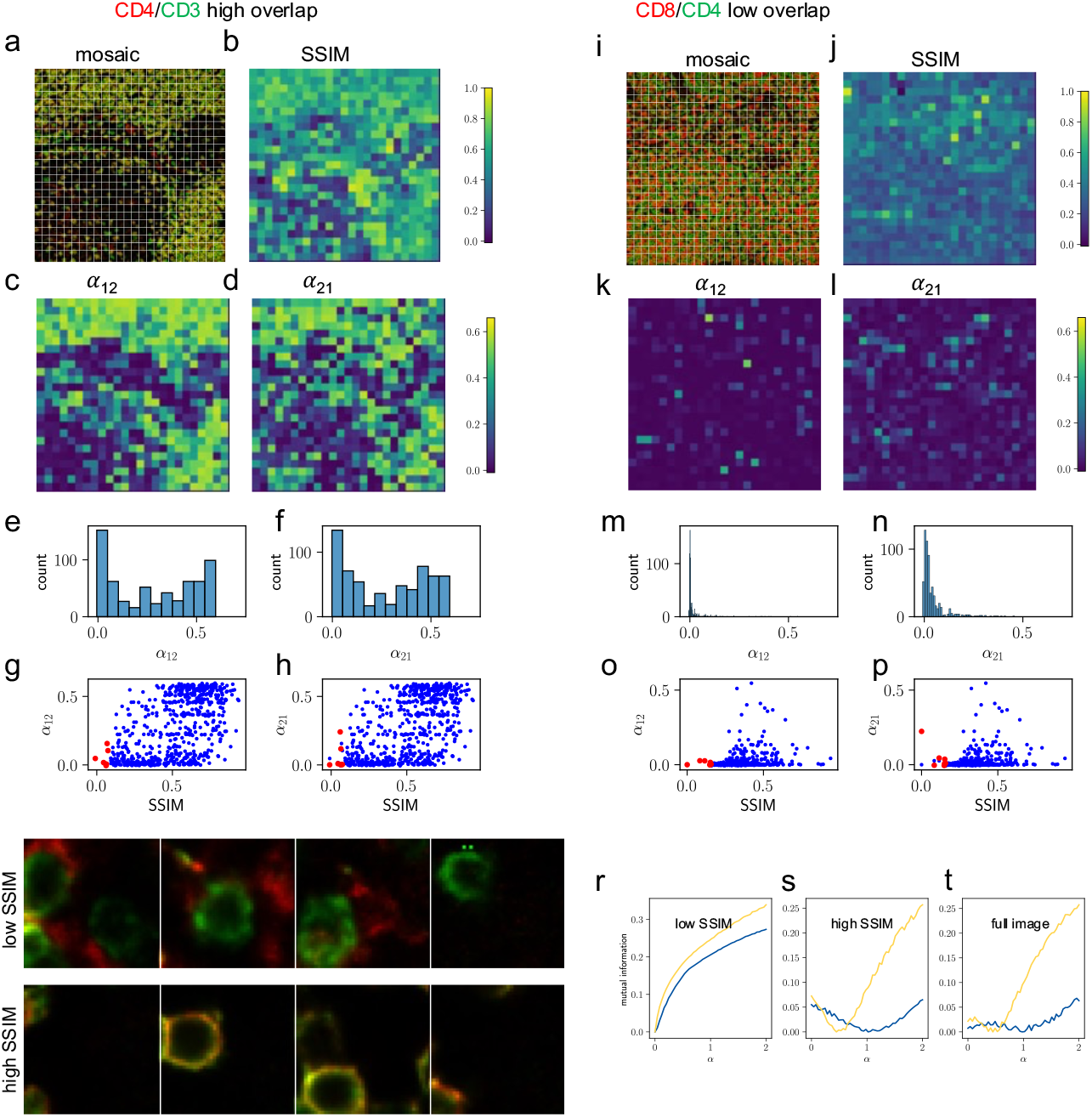
Mosaic PICASSO. (a) The two-color image of CD4/CD3 is divided into mosaics. (b) Spatial Structural Similarity Index (SSIM) is calculated for each mosaic. (c-d) Application of PICASSO to each mosaic recovers the mixing ratios *α*_12_ and *α*_21_, which range from 0 to 0.6 across mosaics, as shown in the histograms (e-f). (g-h) 2D scatter plot of the PICASSO-recovered cross-talk coefficients *α*_12_ and *α*_21_ for each mosaic plotted against the corresponding SSIM. The red dots represent the mosaics in the lower 1st percentile of SSIM. For comparison, (i-p) low overlap targets (CD8/CD4) exhibit smaller spatial heterogeneity in *α*_12_ and *α*_21_. (q) Representative mosaics with lower 1^st^ percentile SSIM (top) and in the upper 50th percentile SSIM (bottom) from the CD4/CD3 images are shown. (r-t) MI curves calculated from the mosaics in the lower 1st percentile of SSIM binned together (r) and the mosaics in the upper 50th percentile of SSIM (s), compared to the full CD4/CD3 image (t).

The estimated mixing ratios α_12_ and α_21_ for each mosaic are depicted in Fig. 2c and d, respectively. The histograms of these ratios demonstrate significant variations ranging from 0 to 0.8 across different mosaics (Fig. 2e, f), highlighting the pronounced spatial heterogeneity. This finding emphasizes that selecting different ROIs within an image can lead to varying unmixing results.

For comparison, we performed the same mosaic analysis on an image expected to have low spatial overlap, CD8/CD4 (Fig. 2i-p). The histogram of the estimated mixing ratios shows significantly less heterogeneity compared to the CD4/CD3e image (Fig. 2m, n).

These results demonstrate that mosaic analysis can serve as a quality check for spectroscopic unmixing using mutual information. The presence of significant spatial heterogeneity in the unmixing results suggests potential of overcorrections.

### Mosaic PICASSO reduces overcorrection artifacts

Recognizing the high heterogeneity often seen in tissue images leads us to hypothesize that the spatial overlap of different fluorophores could vary substantially within a single tissue sample. Consequently, utilizing regions of low similarity to estimate the mixing parameters, then applying them to the full region could mitigate the overcorrection that may occur when PICASSO is applied to the entire image. To investigate this, we plotted a scatter plot of the mixing ratios (α_12_ and α_21_) against the SSIM values of each mosaic of the CD4/CD3e image (Fig. 2g, h). Our findings reveal that mosaics with low SSIM values (SSIM < 0.5) predominantly exhibit low mixing ratios near zero, which closely aligns with the ground truth. In contrast, mosaics with high SSIM values (SSIM > 0.5) exhibit a second population with higher and incorrect mixing ratios (α_12_ and α_21_ > 0.5). The analysis suggests that properly selecting sub-regions or mosaics of an image for mutual information-based unmixing can help reduce overcorrection artifacts.

Due to the inherent variations present in biological samples, establishing a universal threshold for defining low similarity regions is impractical. In our study, we adopt a different strategy by identifying mosaics with the lowest Structural Similarity Index (SSIM) as outliers. These exceptional mosaics demonstrate significant deviations from the average, making them representative of extreme cases. This method can be seen as an optimal solution, allowing for the extraction of maximal information to accurately identify mixing without relying on prior knowledge of the samples or reference spectra. Traditional outlier detection methods, such as the Dixon test^10^ and Grubbs test^11^, rely on assumptions of normal distribution that may not hold true for diverse biological samples. Alternatively, the use of a percentile-based threshold can provide a more robust approach^12^. We hypothesize that selecting the mosaics with SSIM values below the 1st percentile can lead to improved unmixing results.

The mosaics belonging to the bottom 1 percentile are highlighted as red dots in Fig 2g, h, and Fig. 2o, p. Their mixing ratios closely approximate the ground truth value of 0 in both high and low overlap images. Fig. 2q displays representative mosaics from the CD4/CD3 images, with the top panels showing mosaics from the lower 1st percentile SSIM and the bottom panels displaying mosaics from the upper 50th percentile SSIM. In the low SSIM panels, the red and green features are clearly separated, whereas in the high SSIM panels, they exhibit significant overlap.

We further analyze the low SSIM (<1st percentile) and high SSIM (>50th percentile) mosaics by combining them into two separate images and applying PICASSO individually. As shown in Fig. 2r, the MI curves obtained from the low SSIM mosaics accurately predict zero mixing, whereas the MI curves from the high SSIM mosaics incorrectly indicate positive mixing (Fig. 2s), similar to the results obtained from the full image analysis (Fig. 2t).

The above simulation suggests that applying PICASSO to low SSIM score regions of an image could yield a much more accurate estimation of the mixing parameters than when applied to the full image. To further test this result, we generated a series of simulated images with varying mixing ratios and tested if Mosaic PICASSO could accurately recover the mixing parameters (Fig. 3a).

**Fig. 3.**
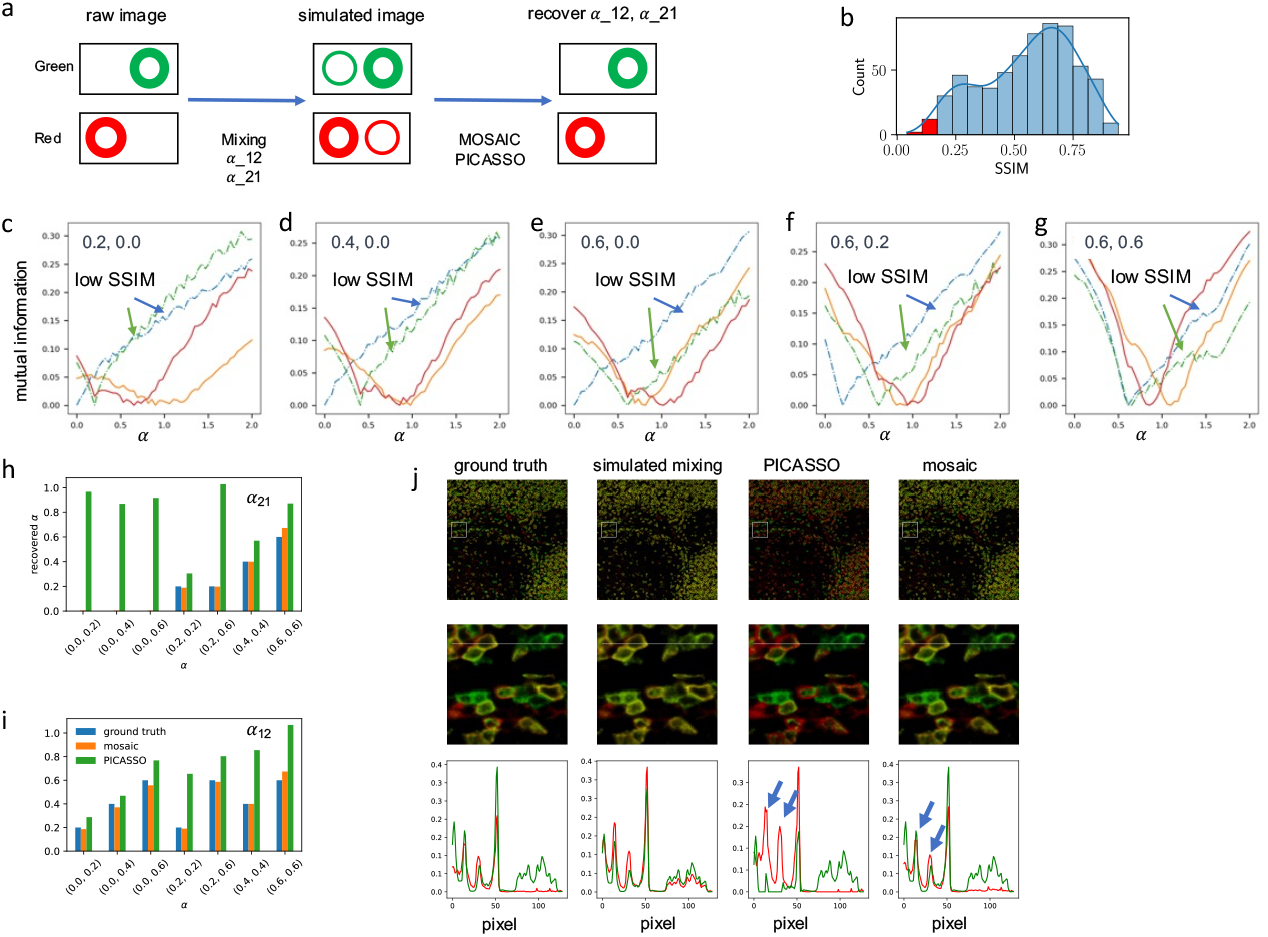
Testing mosaic PICASSO. a) To evaluate Mosaic PICASSO, we simulate spectroscopic mixing of two-color images with varying mixing ratios, *α*_12_ and *α*_21_. We compare the performance of PICASSO with and without the mosaic approach. (b) In the mosaic approach, only the mosaics in the lower 1st percentile of SSIM are used to calculate the mixing ratios. (c-g) The MI versus *α*_12_ and *α*_21_ curves, obtained using Mosaic PICASSO (dash-dot line) and standard PICASSO (solid line), demonstrate that Mosaic PICASSO accurately recovers the mixing ratios, while standard PICASSO significantly overestimates them. The ground truth values for *α*_12_ and *α*_21_ are labeled in each panel. (h-i) Bar plots comparing the ground truth (blue), PICASSO-recovered (green), and Mosaic PICASSO-recovered (yellow) values of *α*_12_ and *α*_21_ show that Mosaic PICASSO exhibits lower overcorrection. (j) The image panels display the ground truth, simulated mixing, PICASSO demixed, and Mosaic PICASSO demixed images for a mixing parameter of (0.2, 0.6). The blue arrows highlight the overcorrection artifacts introduced by PICASSO.

We then apply the mosaic approach to estimate the spectroscopic mixing. For each image, we select the mosaics whose SSIM value is below the 1st percentile and combine them together to calculate the MI curve (Fig. 3b). Fig. 3c-g illustrate the calculated mutual information curves from the low MI mosaics (dash-dot lines) and the full images (solid lines) for various simulated mixed images. The ground truth of the mixing parameters is labeled on each panel. The α value at the minimum of the MI curves represents the calculated mixing parameters, which are shown in the bar plots (Fig. 2h, i). It is evident that Mosaic PICASSO (yellow in Fig. 3h and i) provides significantly more accurate estimation of the mixing parameters (blue in Fig. 3h, i) compared to the original PICASSO (green in Fig. 3h and i) in every case.

The cross-section profile in Fig. 3j shows that PICASSO significantly overcorrects (highlighted by blue arrows), resulting in the green and red channels becoming uncorrelated. In comparison, mosaic PICASSO maximally preserves the spatial correlation and removes the spectroscopic mixing as well, yielding results that are nearly the same as the ground truth.

### The Mosaic method is compatible to multiplex images

PICASSO can be expanded to *n* fluorphores^7^ via an iterative optimization procedure. 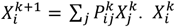 denotes the image of the *i*-th channel after *k* iterations. 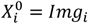 Here *Img*_*i*_ is the unmixed *i*-th channel image. The 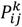 matrix is updated every step as the following:

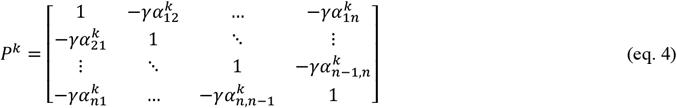

where the γ denotes update step size, and 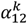 is calculated by equation 3.

We have integrated the mosaic step into the iterative PICASSO algorithm (Fig. 4a). During each iteration step, 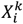 and 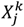, which represents the *i*-th and *j-*th channel images after the *k*-th iteration, are segmented into mosaics of two-color channels. Both MI and SSIM are calculated for each mosaic. The group of mosaics with lower SSIM (<1 percentile) scores is selected for the calculation of the mixing parameters, *α*. These steps are identical to those used in the 2-color case. In the 3-color case, these processes are repeated independently for each pair of two channels: (Img1, Img2), (Img1, Img3), and (Img2, Img3). Each pair has its own group of mosaics selected independently to generate *α* values. These *α* values are then used to generate the *P* matrix, which is subsequently utilized to update all the images.

**Fig. 4.**
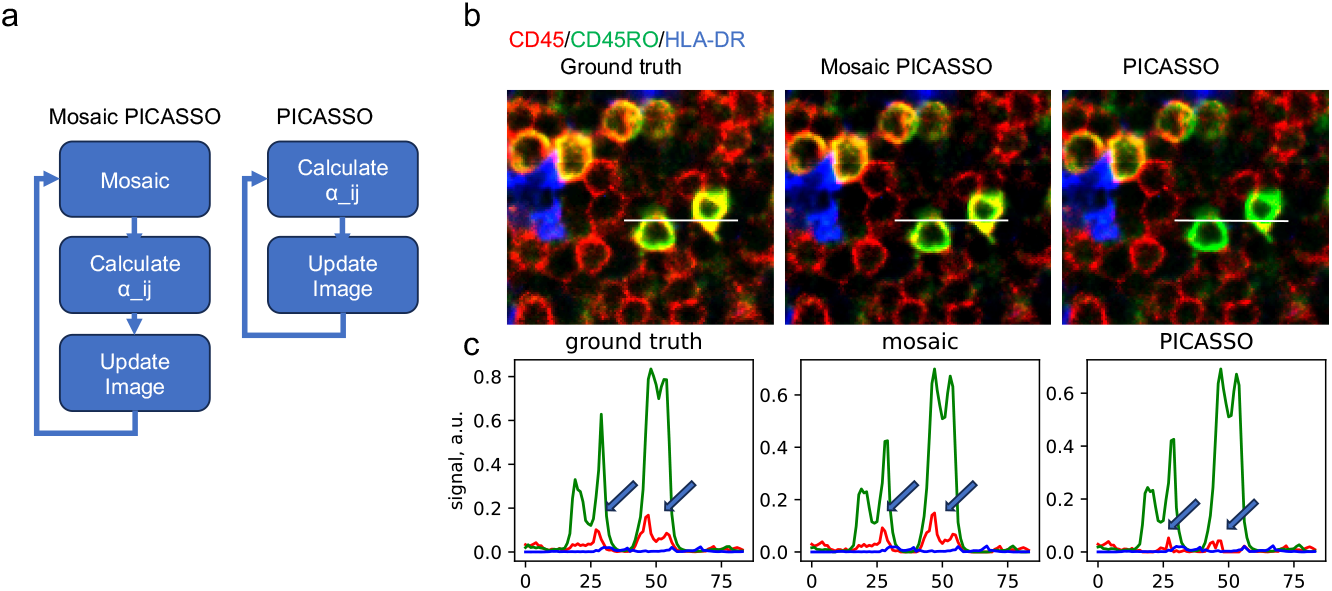
Assessing mosaic PICASSO on 3-plex images. a) The mosaic approach can be seamlessly integrated into the iterative unmixing process of PICASSO and can be extended to handle three or more color images. (b) The panels demonstrate the unmixing results of a 3-color image using PICASSO and Mosaic PICASSO. (c) The profile plot illustrates that Mosaic PICASSO achieves higher accuracy compared to PICASSO.

Fig. 4b demonstrates a 3-plex image of CD45/CD45RO/HLA-DR. The image is taken from a previous Cy-CIF dataset with zero spectral mixing. In an ideal scenario, the three-color image should exhibit zero spectroscopic mixing. As anticipated, Mosaic PICASSO accurately unmixed the three channels, while the original PICASSO significantly overcorrected. The cross-sectional profile illustrates that the original PICASSO nearly eradicated the CD45 channel, greatly reducing the spatial correlation between CD45 and CD45RO. These results highlight the limitations of utilizing MI for spatial biology and reaffirm the effectiveness of implementing Mosaic PICASSO to overcome these challenges.

## Discussion

Our study highlights a limitation of mutual information-based techniques for spectral unmixing when dealing with significant spatial co-localization. The original PICASSO algorithm, which relies on mutual information as a measure of channel dependence, struggles to distinguish between spectral mixing and spatial co-localization effectively. This suggests that mutual information-based methods may not fully capture the complexity of spatial co-localization in biological samples. In response to this challenge, we propose the Mosaic PICASSO approach, which leverages regions of low similarity to estimate spectroscopic overlap. Importantly, the mosaic step can be seamlessly integrated into the iterative PICASSO process, resulting in improved performance compared to the conventional PICASSO approach that uses entire images. Overall, PICASSO represents a significant advancement in ultra-multiplex fluorescence imaging by enabling blind demixing and eliminating the need for costly and error-prone spectroscopic calibration steps. The introduction of the mosaic approach further enhances the accuracy of PICASSO, particularly in samples with diverse levels of mixed spatially colocalized targets confounded by spectral mixing, opening doors to new possibilities in the field of ultra-multiplexed fluorescence imaging.

## Methods

### Normalized mutual information calculation

We first review the definition of mutual information as it pertains to two image channels. For channels Img1 and Img2, the normalized Mutual Information (MI) is calculated using the following formula:

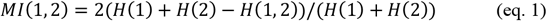

Here, H(1) is the entropy of image 1, H(2) is the entropy of image 2, and H(1, 2) is the joint entropy of images 1 and 2. The entropy H of an image could be calculated from the pixel intensity as:

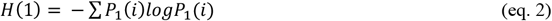

In this equation, ‘i’ represents the pixel intensity in the image, and *P*_1_(*i*) is the probability of occurrence of a pixel of intensity ‘i’ in image 1. This probability can be determined from an intensity histogram of image 1. H(1, 2) represents the joint entropy of the two images, and can be calculated as follows:

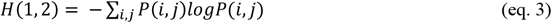

In this case, *P*(*i,j*) is the joint probability of occurrence of a pixel of intensity ‘i’ in image 1 and intensity ‘j’ in image 2. This joint probability can be determined from a 2-dimensional intensity histogram of image 1 and 2.

### SSIM and Pearson’s correlation coefficients

The Structural Similarity Index (SSIM) is computed using the ‘structural_similarity’ function from the ‘skimage.metrics’ module in the Python package ‘scikit-image’. The Pearson’s correlation coefficient is calculated using the ‘pearsonr’ function from the ‘scipy.stats’ module.

### PICASSO

The original PICASSO is a MATLAB package. However, we have implemented the same PICASSO algorithm in Python for our study.

### Mosaic PICASSO

A mosaic module has been implemented in Python for our study. In the case of multiplex images, the mosaic step is incorporated into each iterative round for any pair of channels. In each round, the module divides an image of the two channels into multiple mosaics, with each mosaic being chosen to be approximately the size of 1 to 2 cell diameters. SSIM scores are calculated for each mosaic, and then the mosaics with SSIM scores below the 1st percentile are combined together to calculate the mixing parameters between the two channels. The obtained mixing parameters are used to generate a mixing matrix *P*, which is then used to update the entire image.

### Code

The code is available upon request.

---

**Supplementary Fig. 1.**
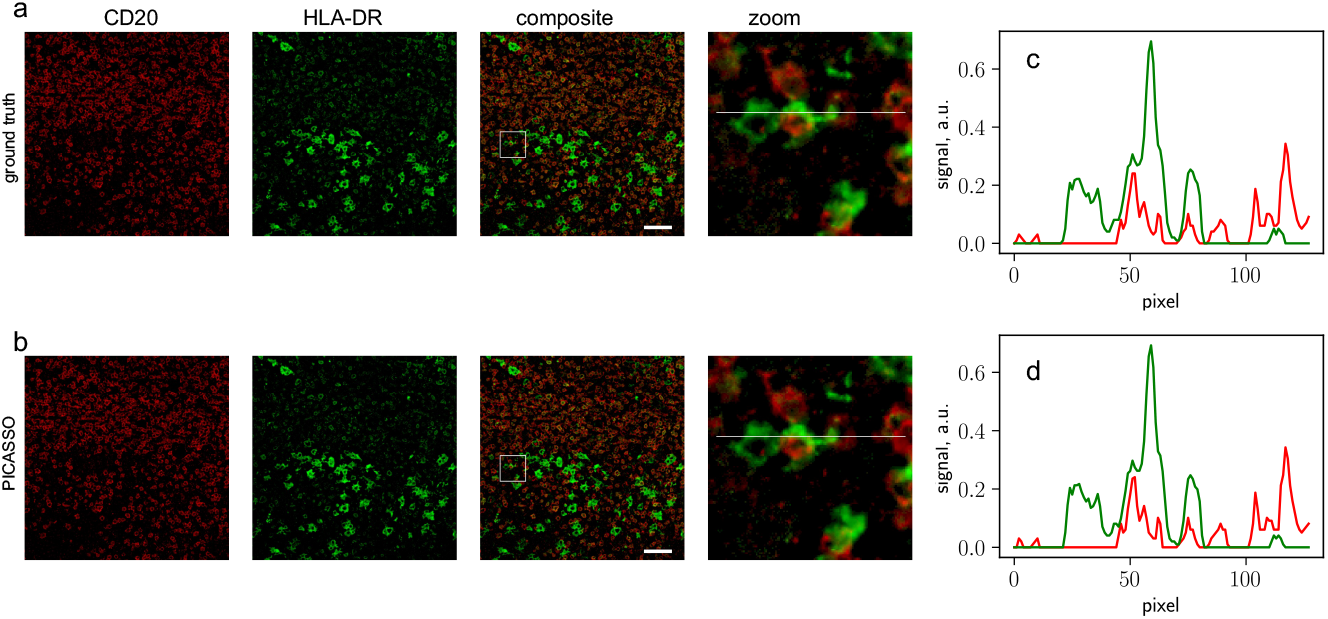
(a) and (b) PICASSO correctly unmix the low overlap target pairs CD20/HLA-DR. The profiles of the ground truth (c) and PICASSO unmixing results (d) are nearly identical.

**Supplementary Fig. 2.**
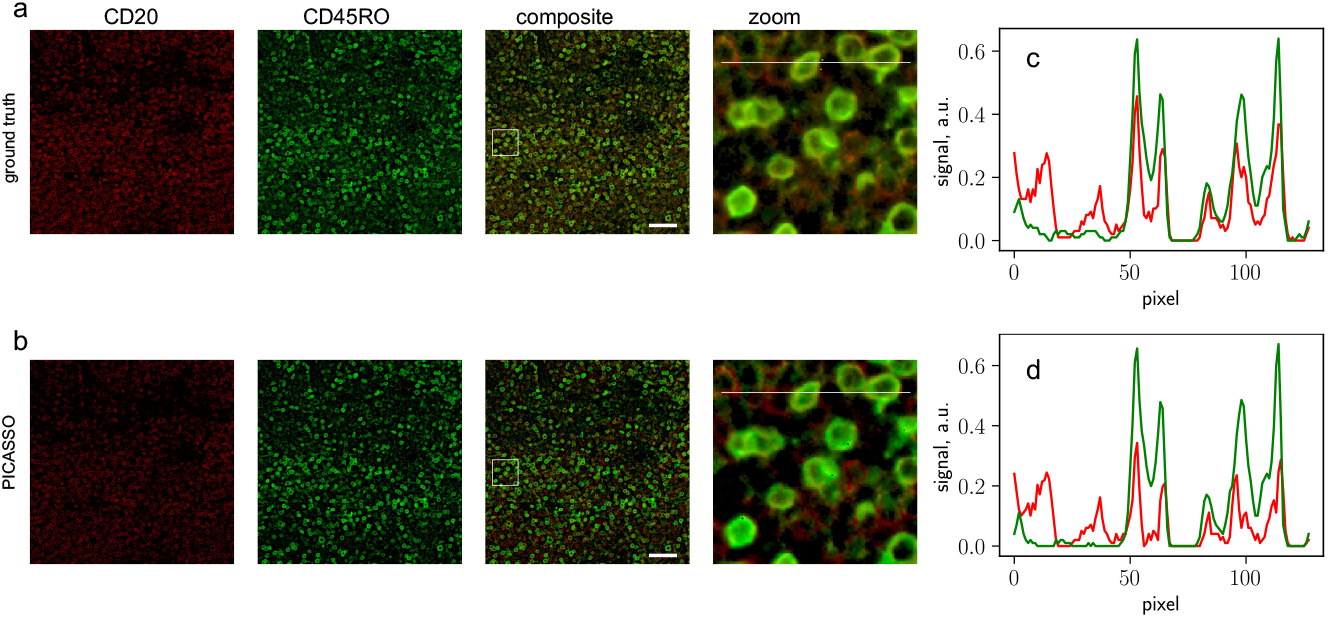
(a) and (b) PICASSO overcorrects for the overlap target pairs CD20/CD45RO. Comparing the profiles of the ground truth (c) and PICASSO unmixing results (d) reveals that the red channel is overly subtracted after demixing.

